# Reprogrammable 4D Tissue Engineering Hydrogel Scaffold via Reversible Ion Printing

**DOI:** 10.1101/2025.02.11.637741

**Authors:** Aixiang Ding, Fang Tang, Eben Alsberg

## Abstract

Shape changeable hydrogel scaffolds recapitulating morphological dynamism of native tissues have emerged as an elegant tool for future tissue engineering (TE) applications, due to their capability to create morphodynamical tissues with complex architectures. Hydrogel scaffolds capable of preprogrammable, reprogrammable and/or reversible shape transformations would widely expand the scope of possible temporal shape changes. Current morphable hydrogels are mostly based on multimaterial, multilayered structures, which involve complicated and time-consuming fabrication protocols, and are often limited to single unidirectional deformation. This work reports on the development of a transformable hydrogel system using a fast, simple, and robust fabrication approach for manipulating the shapes of soft tissues at defined maturation states. Simply by using an ion-transfer printing (ITP) technology (i.e., transferring Ca^2+^ from an ion reservoir with filter paper and subsequent covering on a preformed alginate-derived hydrogel), a tunable Ca^2+^ crosslinking density gradient across the hydrogel thickness has been incorporated, which enables preprogrammable deformations upon further swelling in cell culture media. Combining with a surface patterning technology, cell-laden constructs (bioconstructs) capable of morphing in multiple directions are deformed into sophisticated configurations. Not only can the deformed bioconstructs recover their original shapes by chemical treatment, but at user-defined times they can also be incorporated with new, different spatially controlled gradient crosslinking via the ITP process, conferring 3D bioconstruct shape reprogrammability. In this manner, unique “3D-to-3D” shape conversions have been realized. Finally, we demonstrated effective shape manipulation in engineered cartilage-like tissue constructs using this strategy. These morphable scaffolds may advance 4D TE by enabling more sophisticated spatiotemporal control over construct shape evolution.

## Introduction

Living tissues during development can change their shape to form final functional tissue architecture, better interact with neighboring tissues and/or achieve better nutrient and waste exchange with the surrounding environment. A representative example of morphologically dynamic tissue development is branching morphogenesis whereby the tissues grow into tree-like architectures through evagination of epithelial tubes to maximize the surface area.^1^ Therefore, to biomimic aspects of tissue maturation with tissue engineering (TE), which sometimes utilizes isolated cells and biodegradable scaffolds to produce artificial equivalents for patients with diseased or damaged tissues^2^, it may be valuable to not only regulate cell proliferation, cell differentiation, and secretion of extracellular matrix (ECM) but also potentially closely imitate the dynamic architectural form changes that occur during development and responses of tissues to native stimuli.^3^ However, traditional TE strategies mostly focus on tuning scaffold macro- and microstructures that resembled mature tissue architectures and often ignore the developmental morphodynamics of native tissues.^4^ Since continuous shape evolution occurs during *in vivo* maturation, new approaches have focused on synthesizing shape transformable (termed 4D) scaffolds to enable 4D biomimetic TE.

Hydrogels synthesized from the crosslinking of hydrophilic polymers and macromers^5^ as scaffolding materials have garnered prominence in TE since they are readily available and can be engineered to resemble natural tissue ECM both physically and biochemically.^6^ Geometrically programmed hydrogels with the potential to achieve shape transformations are highly desirable for recapitulating tissue morphodynamics and, as such, work toward this goal has facilitated the development of shape-morphing TE hydrogel scaffolds.^7–8^ The engagement of shape transformability breaches the boundary of conventional geometrically inert 3D TE scaffolds and is considered an important next-generation TE technology.^9–10^ In addition to the dynamically adjustable architectures, the employment of geometrically transforming hydrogels also presents the potential to fabricate TE scaffolds with more sophisticated structures compared to traditional inanimate hydrogels.^11–12^

Strain mismatch acting as the driving force to trigger the shape changes is a primary strategy for designing transformable hydrogel scaffolds.^13^ In this regard, transformable hydrogel scaffolds are often devised as multilayers comprised of materials with different swelling properties in different layers.^14–16^ Nevertheless, multilayered hydrogels are structurally complex, necessitate multistep preparation, and may encounter delamination issues due to insufficient interfacial adhesion.^17^ In contrast, single-layer hydrogel scaffolds are simple in structure and fabrication process but need careful anisotropy incorporation into the structures by controlling the spatial arrangement of the materials used. Therefore, the selection of smart materials with defined responsiveness and the structural design exerts a crucial impact on the reconfigurability capacity of final scaffold products. In this respect, natural polymers such as alginate, hyaluronic acid, silk, gelatin, and chitosan and synthetic polymers such as polyethylene glycol (PEG) represent potential base materials for 4D systems,^18^ with alginate being of particular interest as the shape of cell-laden hydrogels composed of this material can respond to divalent ions as a physiological stimulus.^19^ However, to induce shape morphing, alginate hydrogels usually need to be formed with other materials to generate structural heterogeneity and/or be cast carefully into a single layer with a crosslinking density gradient.^20^ Hence, the fabrication of a responsive single-layered hydrogel scaffold often involves time-consuming protocols such as slow phase diffusion between materials^21^ or complicated photocuring procedures^22–24^. Moreover, reported single-layered alginate hydrogels typically can only undergo a single unidirectional shape morphing,^25^ which limits their application in practical scenarios wherein tissues should exhibit complicated developmental dynamism involving multiple different shape changes over time.

Herein, we present a facile, fast fabrication strategy to produce single-layered one-material hydrogel scaffolds with reversible, reprogrammable shape-morphing performance. Oxidized and methacrylated alginate (OMA) hydrogel constructs encapsulating living cells were programmed to perform prescribed shape transformations. Photocured cell-laden OMA constructs (bioconstructs) were physically crosslinked via an ion-transfer printing (ITP) approach (Scheme 1) to form an ionic crosslinking gradient throughout the hydrogel thickness, due to the decreasing ion concentration along the diffusion direction,^26^ with the top surface directly exposed to the Ca^2+^-soaked filter paper forming high-crosslinking density and tight polymer networks and the bottom surface crosslinked by the diffused Ca^2+^ forming low-crosslinking density and loose polymer networks. The as-prepared large gradient bioconstructs were subsequently tailored into specific geometries to initiate deformation upon culture in cell growth medium (GM) to generate predefined shapes. Furthermore, removing the crosslinker Ca^2+^ ion by a stronger chelator such as ethylenediaminetetraacetic acid (EDTA) at a cytocompatible concentration resulted in internal stress release while maintaining high cell survival. As a result, the bent bioconstructs were straightened and could be reprogrammed by ITP to undergo a different deformation process (*e.g.*, opposite-direction bending). With the proposed approach, we demonstrated “3D-to-3D” morphological conversions and for the first time validated its effectiveness to reshape differentiated tissue-like constructs, revealing its capability to program a ready-to-use 3D bioconstruct of interest. Our results suggest the huge potential of this reversible ion printing technology in reprogrammable 4D TE.

**Scheme 1.**
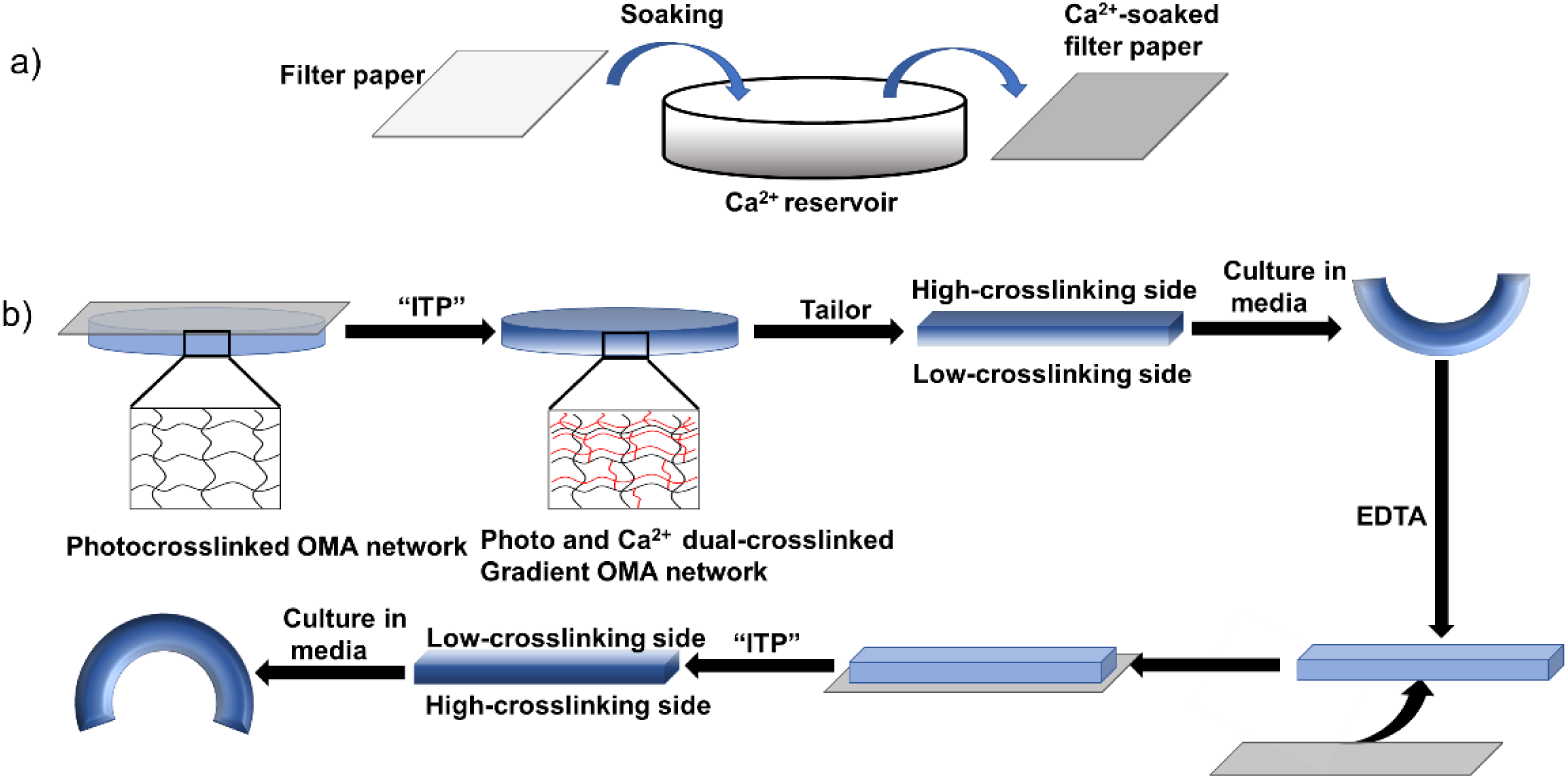
The fabrication of a shape-transformable hydrogel TE scaffold and its shape morphing and reprogramming. (a) Soaking filter paper in Ca^2+^ solution for ions transferring. (b) ITP-resultant crosslinking density gradient in OMA hydrogel scaffold and its subsequent shape transformation upon culture in solution and the reprogramming process.

## Results and discussion

Dual-crosslinkable OMA macromers^27–28^ were synthesized as the main hydrogel material (Scheme S1). The OMA macromers in the presence of a photoinitiator were covalently crosslinked into a stable hydrogel network via photocuring. This was evidenced by the largely weakened methacrylate proton signals (H_a_ and H_b_ in dotted rectangles) in Figure S1 after photoirradiation. The degradation profile of photocured OMA hydrogels plateaued after 1-day culture in GM in an incubator at 37 °C with 5% CO_2_ (Figure S2). This photocured OMA hydrogel can be further ionically crosslinked by Ca^2+^ ions to form dual-crosslinked networks (Scheme S2). Culturing only photocured OMA hydrogels in aqueous solutions such as water (H_2_O), phosphate-buffered saline (PBS) (pH 7.4), and GM induced volumetric expansion (swelling), accompanied by a small range of variation in storage modulus (G’) and a large decrease in Young’s modulus (Figures S3 and S4). In contrast, culturing the hydrogels in 0.5 M Ca^2+^ solution brought about obvious volumetric shrinkage (deswelling), which was accompanied by a large increase in G’ and Young’s modulus. These results confirmed the effective binding of OMAs with Ca^2+^ ions, leading to tightening and toughening of the hydrogel networks. Since the swelling of OMA hydrogels decreases with the increasing polymer network crosslinking density^7^, it was conjectured that the gradient OMA hydrogels resulting from ITP would exhibit out-of-plane deformation with the higher-crosslinking side on the concave side(Scheme S2).

The shape-morphing behavior of the gradient hydrogels strongly relies on the incubation solution, macromer concentrations, Ca^2+^ concentration of ion reservoir, and ITP time. To examine the impact of those parameters on the shape-morphing behaviors, hydrogel bars with dimensions of 15 mm (length) × 2 mm (width) × 1 mm (thickness) were fabricated as a simplified prototype. As can be seen in Figure 1, all those parameters exerted a clear impact on the shape morphing. Generally, parameters contributing to larger swelling and/or greater gradient range across the bar thickness give rise to more pronounced hydrogel deformation. For example, hydrogel bars cultured in H_2_O displayed a significantly higher bending due to the significantly higher swelling (Figure S3b) than those cultured in PBS (pH 7.4) and GM (Figure 1a), while the diminished hydrogel swelling due to the increase in macromer concentration (Figure S5) resulted in a rapid decrease in the bending angle (Figure 1b). Likewise, increasing Ca^2+^ concentration (Figure 1c) and extension of ITP time (Figure 1d) that benefits a larger gradient range led to larger bending angles. The results suggest that the deformation output could be programmed by tuning these parameters. Unless otherwise specified, OMA hydrogels made by 3% w/v macromer concentration, 0.5 M Ca^2+^ reservoir, and 1.0 min ITP time were used for the experiments and results discussed below.

**Figure 1.**
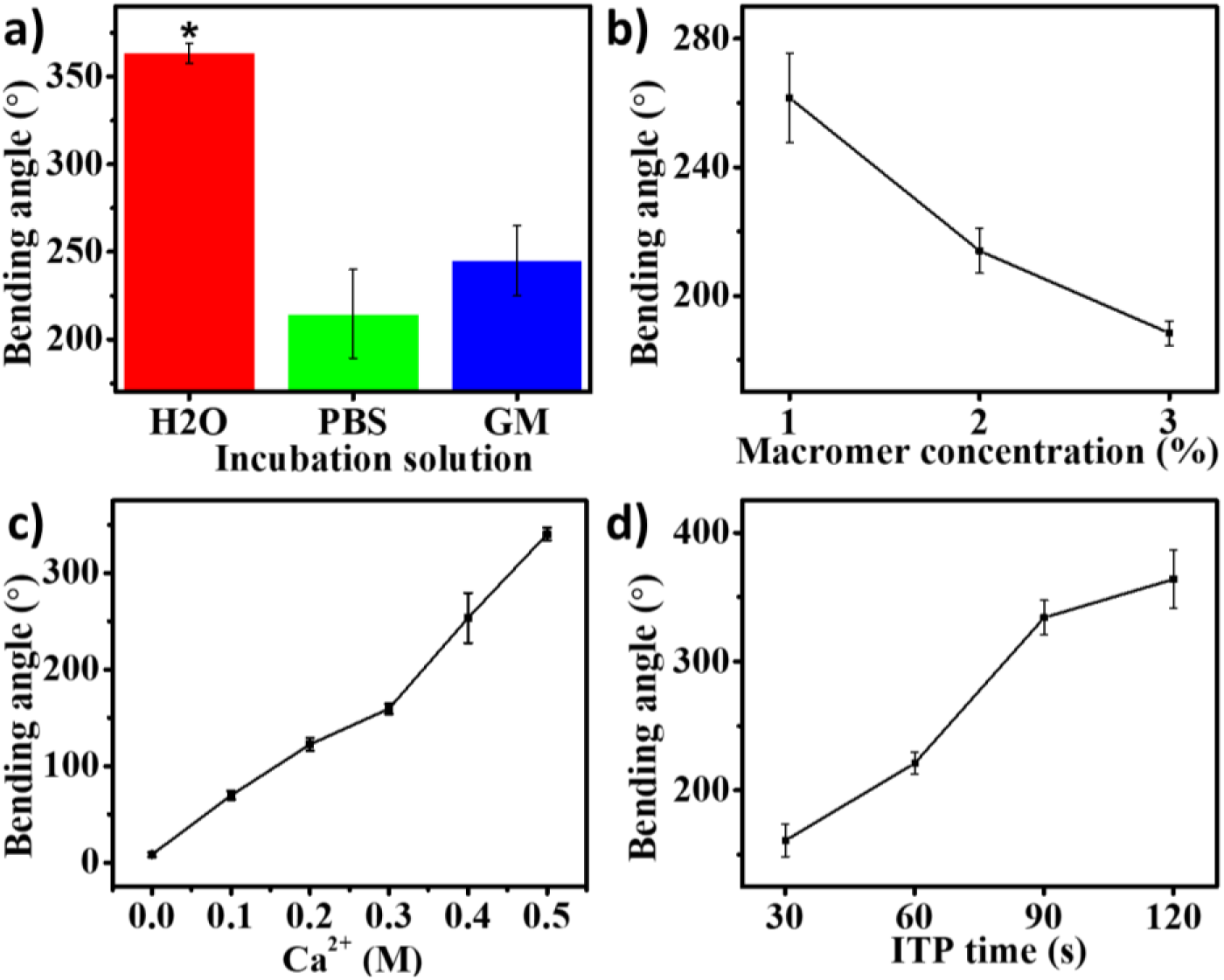
The impact of (a) incubation solution, (b) macromer concentration, (c) Ca^2+^ concentration of ion reservoir, and (d) ITP time on the shape-morphing behaviors of hydrogel bars at room temperature (RT). The other conditions for culture solution study are 3% w/v macromer concentration, 0.5 M Ca^2+^ concentration, and 5 min ITP time; for macromer concentration study are GM as the culture solution, 0.5 M Ca^2+^ concentration, and 5 min ITP time; for Ca^2+^ concentration study are GM as a culture solution, 3% w/v macromer concentration, and 5 min ITP time; and for TIP time study are GM as a culture solution, 3% w/v macromer concentration, and 0.5 M Ca^2+^ concentration. N =3, data are presented as mean ±SD.

Next, gradient OMA hydrogels as cell-laden scaffolds were subjected to incubation in GM to investigate its shape-morphing properties. NIH3T3 cells with a density ranging from 5 to 100 million (M) cells/mL macromer solution were encapsulated and the bending behavior of the resulting hydrogel bars was examined. Similar to the cell-free counterparts, the cell-laden hydrogel bars quickly morphed into a “C” shape in about 3 min in cell culture medium (Figure 2a). There was a downward trend in the bending angle of the cell-laden hydrogel bars with increasing cell density (Figure 2b). With the same method, deformed large bioconstructs were obtained through programming large bioconstructs such as cell-rich hydrogel discs (15 mm diameter × 1.0 mm thickness) and square hydrogel slabs (15 mm length × 1.0 mm thickness) (Figure S6). As expected, the cells within the bent hydrogel bars were presented in round morphology as shown in Figure 2c and maintained high viability indicated by the predominantly green-colored cells in the live/dead staining results in Figure 2d, wherein the live cells were visualized fluorescently with green color by fluorescein diacetate, while dead cells were visualized with red color by ethidium bromide. The cytocompatibility of the shape-morphing system together with its preprogrammable deformability suggests its reliability for use in 4D TE.

**Figure 2.**
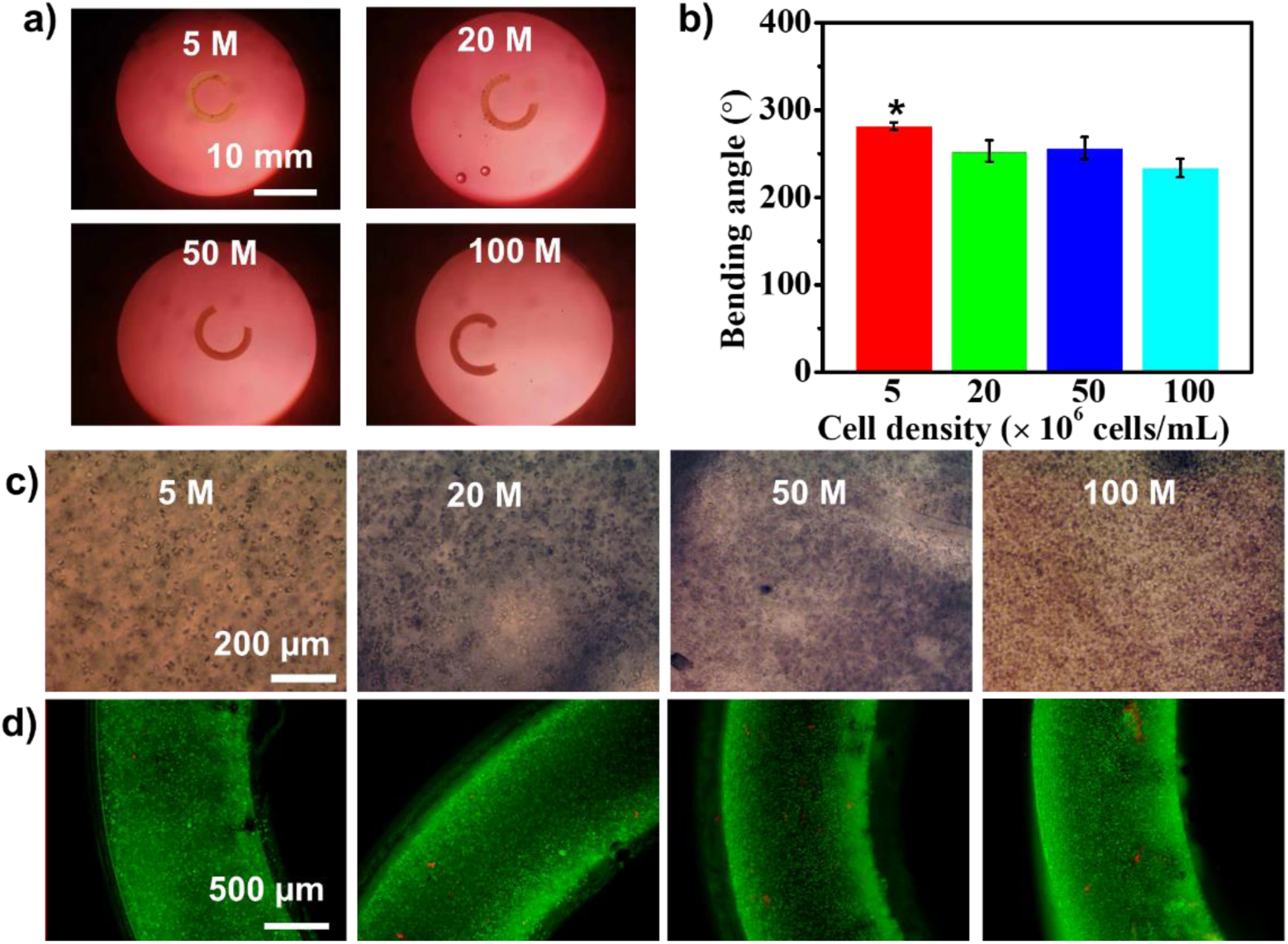
Shape-morphing properties of cell-laden hydrogel bars. (a) Representative photographs show the bent shapes. (b) Quantitative bending angles. (c) Photomicrographs of bright field images. (d) Photomicrographs of representative live/dead fluorescence images. \**p* < 0.05 compared with other groups. N = 3, data are presented as mean ±SD.

The fabrication of cell-laden bioconstructs with complex configurations was explored by harnessing a “patterning ITP” approach. NIH3T3 cells at a density of 20 M/mL hydrogel precursors were used to fabricate bioconstructs unless otherwise stated. In a simple demonstration, we segmentally patterned hydrogel bars to enable asymmetrical, multi-directional bending, yielding “1D-to-2D” shape transformations to afford complex structures with “S” or “S”-like shapes (Figure 3). We next advanced the pattern design to create more complex 3D architectures. Ca^2+^-soaked filter papers with specific geometries were used to create patterned Ca^2+^ crosslinking within disc-shaped bioconstructs (Figure 4). Attributed to the locally suppressed swelling within the Ca^2+^ patterns and the significantly larger swelling in the non-patterned regions, various complex 3D bioconstructs with “wave” or “flower” like structures were obtained via “2D-to-3D” shape transformations after being cultured in GM for 5.0 min. Thus, this simple patterning process was demonstrated to be effective in manipulating the bioconstruct shapes via a set of programmed cooperative deformations.

**Figure 3.**
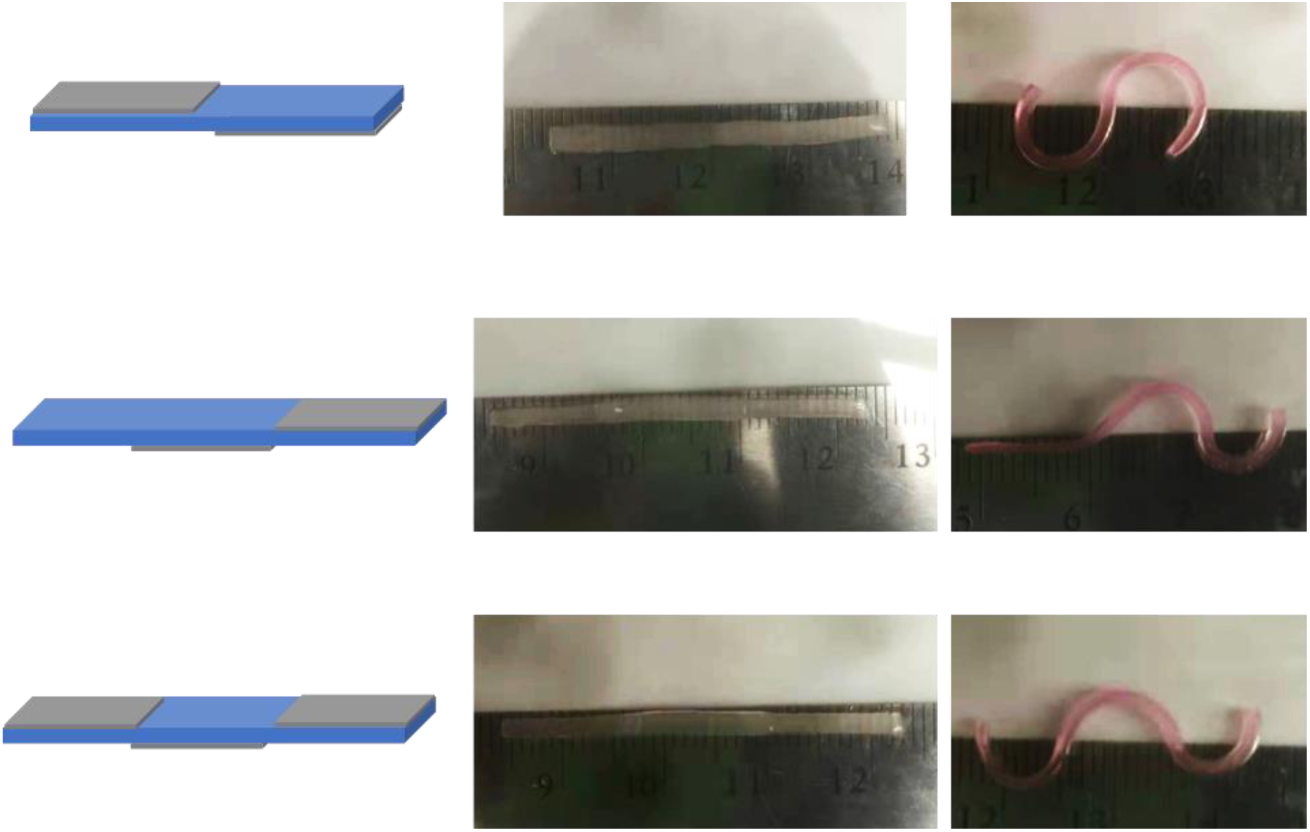
Segmental patterning of cell-laden hydrogel bars and the corresponding resultant multi-directional bending complex 2D bioconstructs. Hydrogel length: 30 mm (upper), 39 mm (middle), and 39 mm (bottom), hydrogel width (2 mm), pattern length: 15 mm.

**Figure 4.**
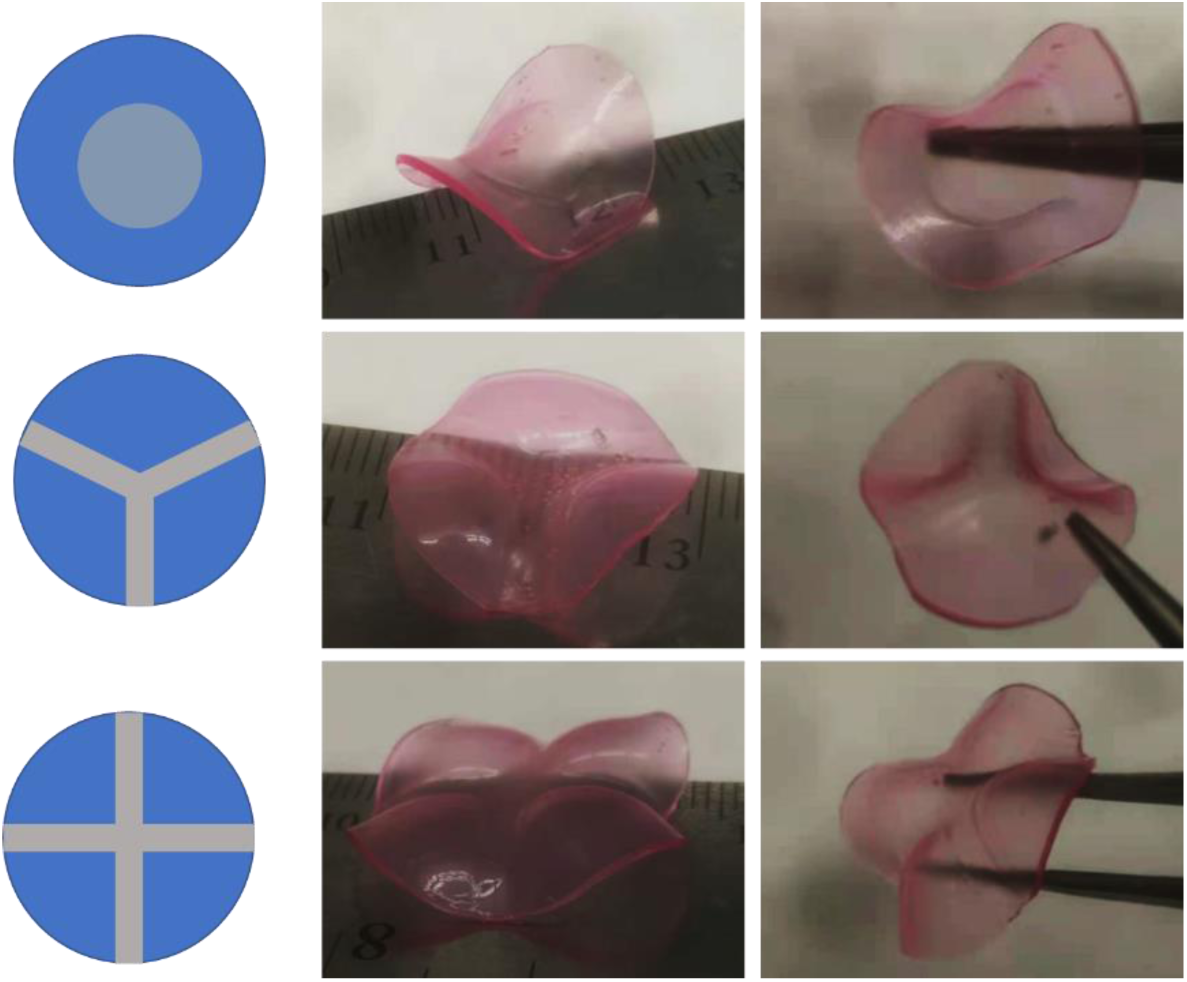
Patterning of cell-laden hydrogel discs and the resulting complex 3D bioconstructs after deformation. Hydrogel diameter: 20 mm, pattern width 0.3 mm.

The anisotropic internal strain caused by the gradient Ca^2+^ crosslinking can be erased by removing the Ca^2+^ from the hydrogel networks with EDTA, which in turn relaxes the bent hydrogel bars and further leads to hydrogel straightening. A subsequent reprogramming via a second ITP process could induce the hydrogel re-bending in a prescribed manner (e.g., bending in the opposite direction). As such, shape reprogramming could be achieved. Figure 5a illustrates a typical reprogramming process. Cell-free hydrogel bars were first employed to verify reprogrammability. As hypothesized, bent hydrogel bars in H_2_O were able to recover to their relaxed states (near-straight shapes) after soaking in EDTA (5 mM) for about 60 min at RT. The straightened hydrogel bars at RT regained their deformed states (“C” shapes) in about 10 min of culture after a second ion printing (Figure S7). The results demonstrated that the shape transformation was reversible. Switching the hydrogel bars back and forth between the two distinct states occurred easily and the response time could be shortened by increasing the incubation temperature to 37 °C (Figure S8).

**Figure 5.**
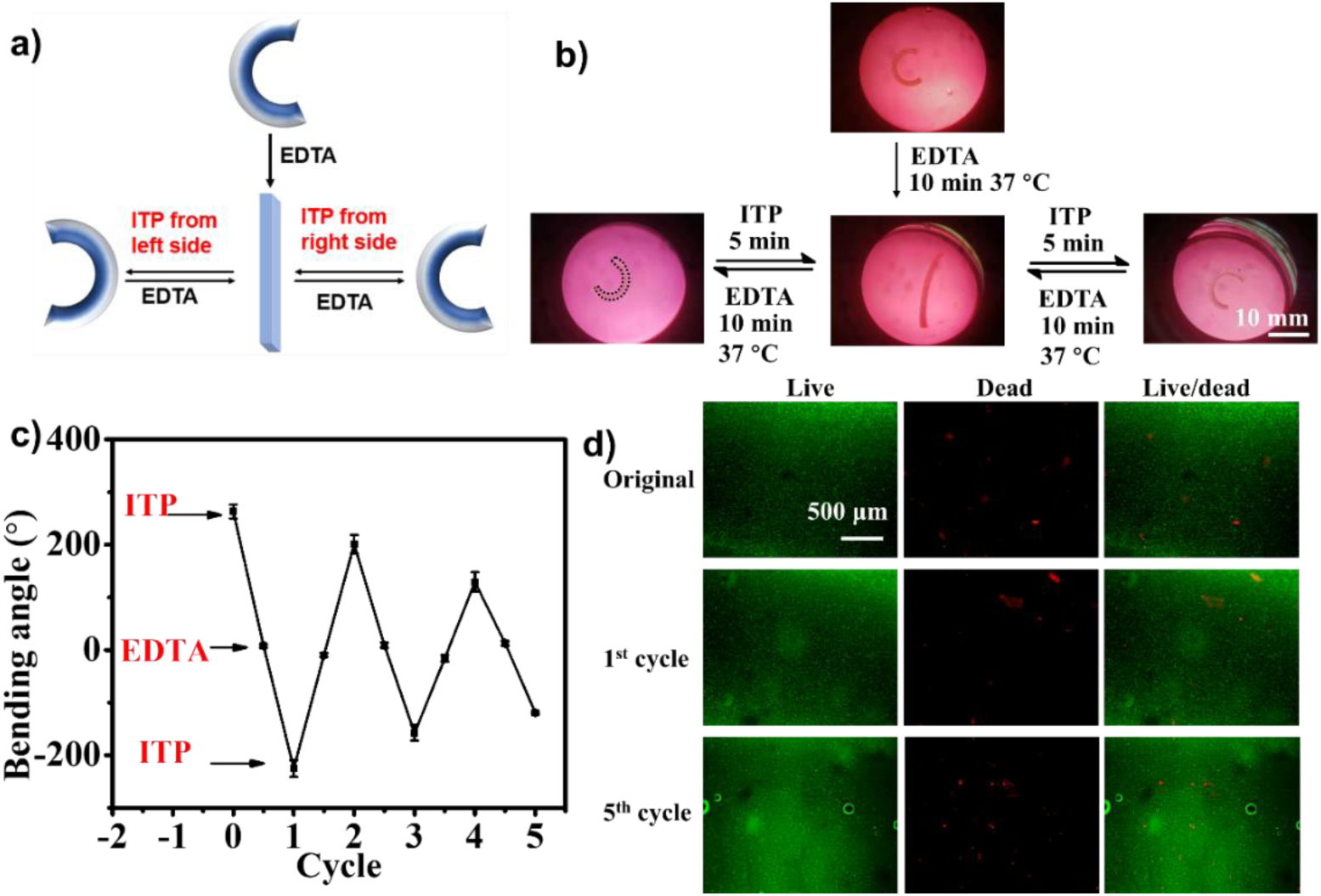
Reprograming of cell-rich hydrogel bars. (a) Schematic illustration of the reprogramming process. (b) Switch of a representative hydrogel bar between stretched and bent states under cell culture conditions. (c) Quantitative bending angles of hydrogel bars at the two states over repetitive cycles. (d) Representative live/dead photomicrographs from different cycles. N =3, data are presented as mean ±SD.

However, signs of fatigue were observed in the following relaxation–reprogramming cycles (Figure S7b), which could be ascribed to the combined effects of the rearrangement of local networks by the repetitive breaking and reforming of the physical bonding^29–30^ and the already swollen networks reducing the range of the resultant gradient crosslinking density. A similar repetitive switching between the relaxed and deformed states by the same reprogramming approach was also observed for cell-rich hydrogel bars (Figures 5b and c), in which opposite-bending programming, quantified by the negative bending angles, was also demonstrated. The encapsulated cells remained highly viable after cycling five times (Figure 5d). Unlike previous work on 4D biofabrication that often demonstrates only “2D-to-3D” shape transformations, unique “3D-to-3D” shape transformations by taking advantage of the reprogrammability were realized in large bioconstructs as shown in Figure 6. In comparison to “2D-to-3D” transformations, the implementation of “3D-to-3D” transformation is deemed much more challenging^31–32^ and may be vaulable in 4D TE for its powerful potential to better mimic the deformations and movements occurring in 3D-structured living organisms.^33^

**Figure 6.**
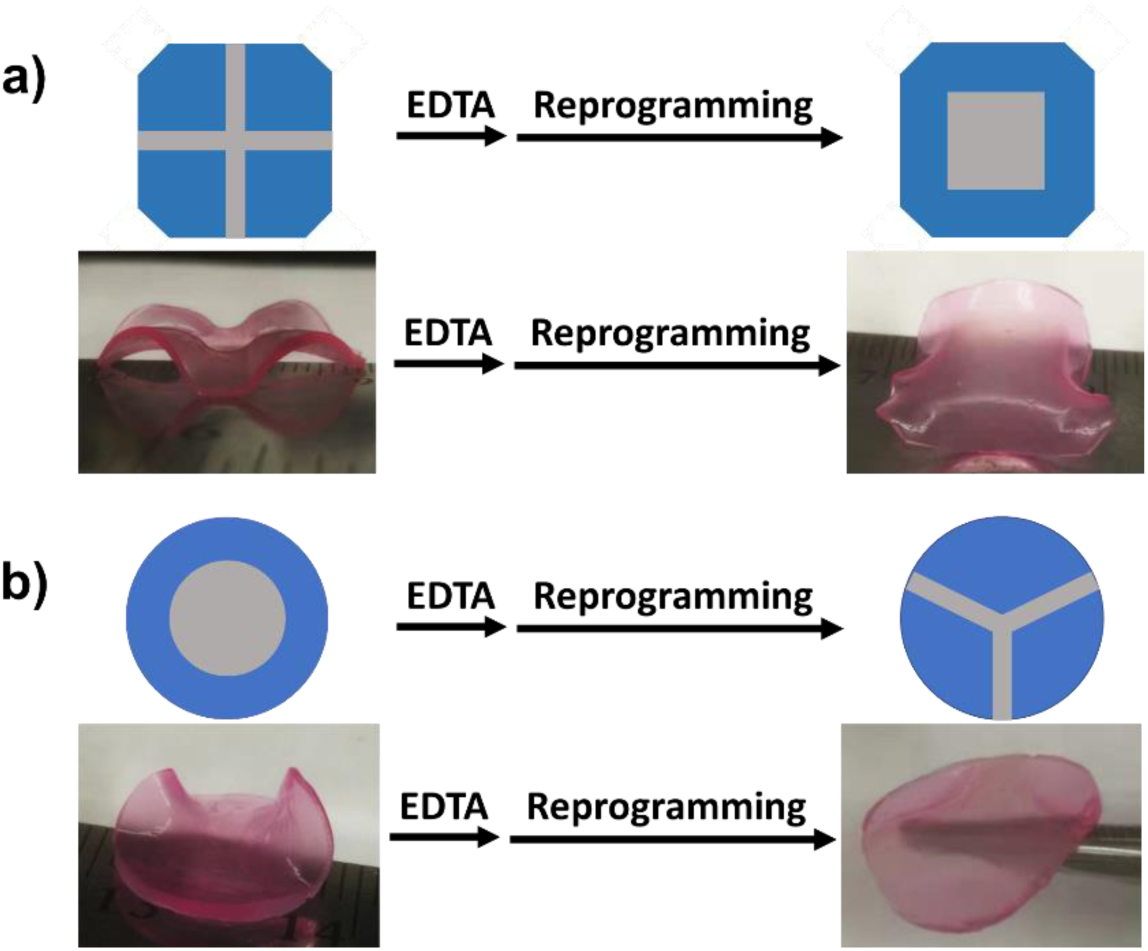
Reprogramming of (a) a hydrogel slab and (b) a hydrogel disc. Length of hydrogel slab: 15 mm; pattern parameter: 3 mm width for cross pattern and 10 mm length for square pattern. Diameter of hydrogel disc: 15 mm; pattern parameter: 10 mm diameter for disc pattern and 3 mm width for “Y” pattern.

The remarkable shape-morphing capability, together with the reprogrammability, inspired us to explore the system’s capacity for driving reprogrammable 4D shape transformation TE. Specifically, 4D chondrogenesis of human mesenchymal stem cells (hMSCs) at a density of 10 M/mL hydrogel precursors was employed as a proof-of-concept model. To promote better cell adhesion to hydrogel networks and better cell survival during long-term culture, 3% (w/v) OMA mixed with 0.5% (w/v) cell-adhesive methacrylated gelatin (GelMA)^34–35^ was used as scaffolding material. The incorporation of a small amount of GelMA did not influence the shape morphing ability of the OMA hydrogels (Figure S9). HMSC-laden bioconstructs in the shape of square slabs were subjected to differentiation in chondrogenic medium over a course of three weeks (21 days), during which the bioconstructs exhibited no morphological conversions but only linear expansions (Figure S10), and cell viability was demonstrated to be high (Figure S11). Biochemical quantification and histological staining were performed to assess the differentiation. The results of the biochemical analysis revealed that, in comparison with the control group (Ctrl, bioconstructs cultured in normal GM), the experimental group (Exp, bioconstructs cultured in differentiation medium) had comparable DNA levels (Figure 7a) but significantly higher glycosaminoglycan (GAG) production, a primary cartilage ECM component (Figures 7b and c). Also, Exp exhibited a significantly higher storage modulus than Ctrl (Figure S12), indicative of the generation of cartilage-like tissue constructs. Histological staining with Hematoxylin and Eosin (H&E) and Safranin O (SafO) showed that the cells on D21 were uniformly distributed throughout the bioconstructs and resided primarily in small pockets surrounded by hydrogel (Figure 7d and e). Moreover, the intense staining of SafO further confirmed substantial GAG production of the engineered cartilage-like tissues. However, the evident SafO staining observed in the Ctrl group may result from nonspecific dye binding to the alginate hydrogels, although the staining intensity was visibly weaker than that of the Exp constructs.

**Figure 7.**
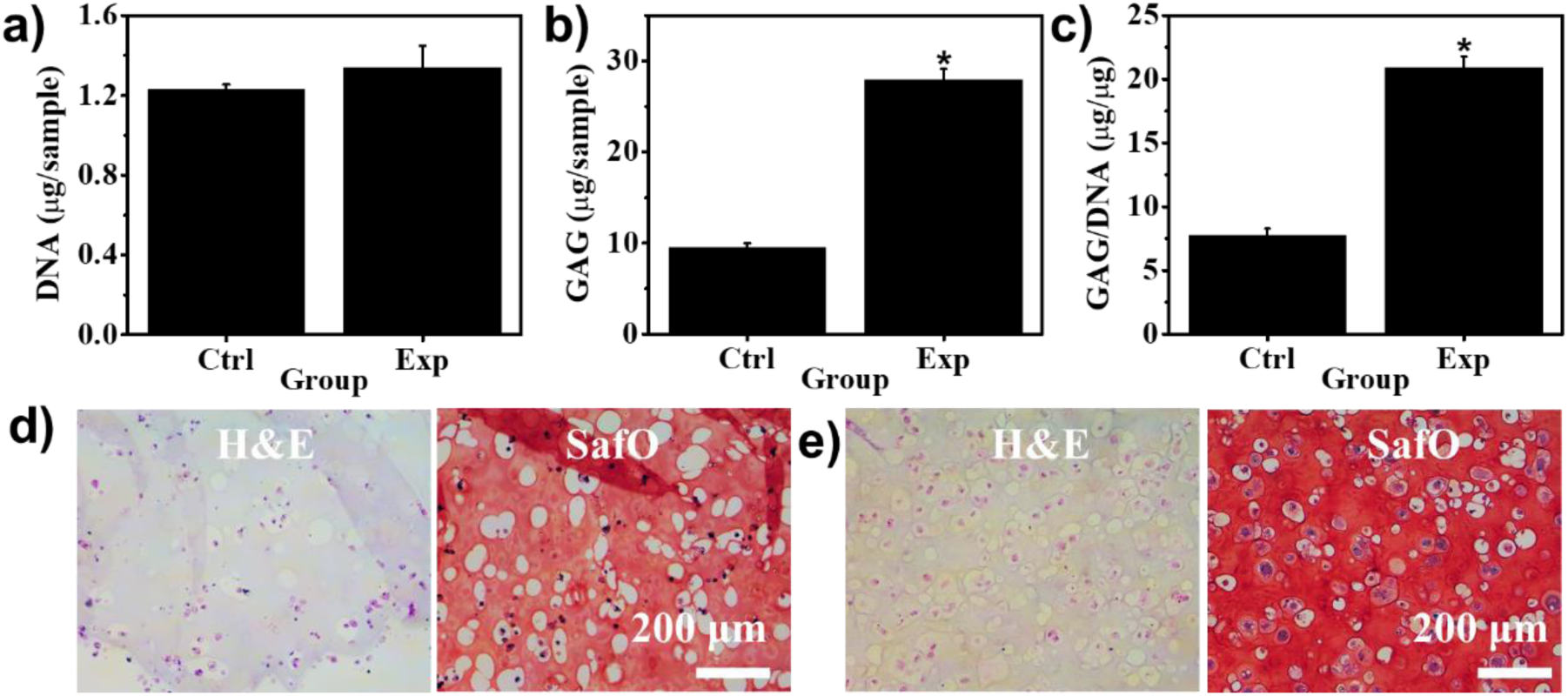
Evaluation of chondrogenesis. Biochemical quantification of (a) DNA levels, (b) GAG production, and (c) GAG/DNA ratios in the control (Ctrl) and experimental (Exp) groups. Photomicrographs of representative H&E and SafO histologically stained samples from (d) the Ctrl group and (e) the Exp group. N =3, data are presented as mean ±SD.

Constructs in different shapes, obtained by cutting the tissue-like specimen, were ion printed to evaluate the shape morphability and reprogrammability of differentiated constructs after 3 weeks of culture in CPM. Like the undifferentiated samples, ion-printed tissue discs and square slabs could also deform into curled geometries (Figure 8a). The results of reprogramming experiments also demonstrated the reprogrammability of these engineered tissues (Figure 8b). However, the deformation extent of these tissue-like bioconstructs was much less pronounced compared to that of undifferentiated bioconstructs, due to the morphing constraints stemming from the extensive stiff ECM produced. Such manipulation of differentiated bioconstructs allows for the construction of functional living tissues with high structural dynamism and complexity, which could be highly valuable for personalized biomedicine for its ability to facilitate seamless integration of engineered tissues into tissue defects of specific patients.^36^

**Figure 8.**
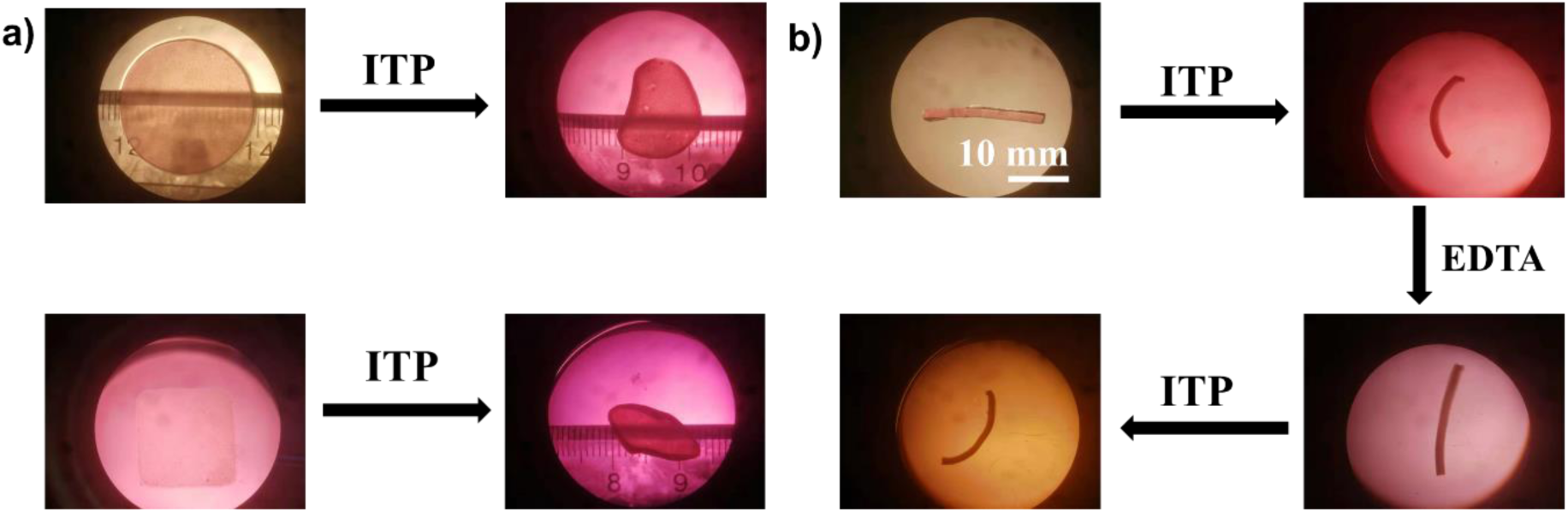
Programming the shape of engineered tissues. (a) Deformation of ion-printed tissue disc and tissue slab. (b) Multiple deformations of a tissue bar via reprogramming.

The focus on biomimicry of native tissue development has recently aroused the need for the development of 4D tissue engineering technologies that can achieve dynamic morphological changes over time.^37–38^ Emerging 4D TE accounts for not only regenerating tissue substitutes with complex conformations but also offers opportunities to imitate the intrinsic dynamism of living tissues^39–41^ and thus may be advantageous in some applications over traditional morphologically static TE, which generally lacks the feature of tissue dynamics^42^. The fast, facile biofabrication approach described herein to establish a reprogrammable scaffold by using inexpensive techniques and simple materials opens a new paradigm for 4D TE applications. Our model provides a robust platform to manipulate construct architectures in multiple ways on demand for both undifferentiated and differentiated bioconstructs at defined times. The shape transformation is driven by the differential swelling caused by the crosslinking density gradient across the hydrogel thickness. We engineered large bioconstructs with sophisticated topologies, such as wavy and flower-shaped structures. High cell survival was detected before and after shape morphing. Impressively, “3D-to-3D” shape transformations have been achieved simply by releasing and rebuilding the internal anisotropic strain. Unprecedently, we demonstrated the potentiality to reprogram the shape of engineered tissues. The developed system outperforms conventional multilayering approaches on 4D biofabrication when it comes to simplicity, efficiency, cytocompatibility, and deformability, and provides morphodynamical means to possibly regulate the biomaterial mechanical properties in a spatiotemporal manner, enhancing the capacity to use scaffolding materials to direct the fate of encapsulated cells.

## Conclusion

On the basis of ITP technology, this work has demonstrated a simple strategy to enable multi-scale structural morphing of soft tissues via physiological relevant stimuli. Preprogrammable and reprogrammable shape morphing of bulk cell-laden alginate hydrogels occur as a result of forming reversible gradient crosslinking. Deliberately tuned parameters including culturing medium, ITP time, macromer concentration, and Ca^2+^ reservoir concentration, in conjugation with specific local crosslinking incorporation by means of surface patterning, allow for controlled, multiple, multi-directional deformations, generating complex tissue constructs at various stages of maturation. Multi-dimensional switching between 1D, 2D, and 3D structures endows the system with excellent biofabrication capabilities, making it adaptable for customized, complex biological and physiological environments. The results collectively presented in our study reveal a reliable platform for 4D TE, which could also be a valuable tool for investigating the role of dynamical architecture in mechanobiology, developmental morphogenesis, bioactuation, and beyond.

## Acknowledgements

The authors gratefully acknowledge funding from the National Institutes of Health’s National Institute of Arthritis and Musculoskeletal and Skin Diseases (R01AR081448) and National Institute of Biomedical Imaging and Bioengineering (R01EB023907), and the Department of Veterans Affairs, Veterans Health Administration, Office of Research and Development, Rehabilitation Research and Development Service (RX004288). The authors also thank Sriramya Ayyagari at University of Illinois at Chicago for reviewing the manuscript. The contents of this publication are solely the responsibility of the authors and do not necessarily represent the official views of the Department of Veterans Affairs or the National Institutes of Health.

## Author contributions

**A. Ding**: conceptualization, methodology, data collection, formal analysis; **F. Tang**: Formal analysis, visualization, writing – original draft; **E. Alsberg**: conceptualization, methodology, formal analysis, resources, supervision, funding acquisition, writing – review & editing.

## Supporting Information

### 1. Experimental

#### 1.1 Chemicals, instruments, and general methods

Unless specified, all solvents and reagents were used without further purification. Sodium alginate (Protanal LF120M, 251 Pa·s) was a generous gift from FMC Biopolymer. Photoinitiator (2-Hydroxy-4’-(2-hydroxyethoxy)-2-methylpropiophenone), fluorescein diacetate (FDA), ethidium bromide (EB), Dulbecco’s Modified Eagle Medium-Low Glucose (DMEM-LG), and fetal bovine serum (FBS) were purchased from Sigma. Insulin transferrin selenium^+^ (ITS^+^) Premix and penicillin/streptomycin (P/S) were purchased from Corning Inc. (Corning, NY). Sodium pyruvate was purchased from HyClone Laboratories (Logan, UT). Non-essential amino acid solution was purchased from Lonza Group (Basel, Switzerland). Ascorbic acid-2-phosphate was purchased from Wako Chemicals USA Inc. (Richmond, VA). Fibroblast growth factor-2 (FGF-2) was purchased from R&D Systems (Minneapolis, MN). Transforming growth factor β1 (TGF-β1) was purchased from PeproTech (Rocky Hill, NJ). *N*-(2-aminoethyl) methacrylate hydrochloride (AEMA) and methacryloxyethyl thiocarbamoyl rhodamine B (RhB) were purchased from Polysciences Inc. (Warrington, PA), and other common chemicals, such as sodium peroxide, methacrylic anhydride, etc., were purchased from Fisher Scientific (Hampton, NH). ^1^H NMR spectra were obtained on a 400 MHz Bruker AVIII HD NMR spectrometer (Billerica, MA) equipped with a 5 mm SmartProbe™ at 25 °C using deuterium oxide (D_2_O) as a solvent and calibrated using (trimethylsilyl)propionic acid-*d*_4_ sodium salt (0.05 w/v %) as an internal reference. DMEM-LG containing 0.05% PI (w/w) was used to dissolve the oxidized methacrylate alginate (OMA) and methacrylate gelatin (GelMA). Cell growth medium (GM) consisted of DMEM-LG with 10% FBS and 1% P/S, and chondrogenic medium consisted of DMEM-LG with 1% ITS^+^ Premix, 100 nM dexamethasone, 1 mM sodium pyruvate, 100 µM non-essential amino acids, 37.5 µg/mL ascorbic acid-2-phosphate and 1% P/S supplemented with 10 ng/mL TGF-β1. Hydrogel images were visualized using a Nikon SMZ-10 Trinocular Stereomicroscope equipped with a digital camera. A microplate reader Molecular Devices iD5 (San Jose, CA) was used to read data from the microplates. A UV device (EXFO OmnicureR S1000-1B, Lumen Dynamics Group, Mississauga, Canada) with an intensity of 18 mW/cm^2^ was used for photocrosslinking. All quantitative data was expressed as mean ±SD. Statistical analysis was performed with one-way analysis of variance (ANOVA) with Tukey honestly significant difference post hoc tests using Origin software (OriginLab Corporation, Northampton, MA). A value of *p* < 0.05 was considered statistically significant.

#### 1.2 Synthesis of OMAs and GelMA

OMA macromers with a theoretical 1% oxidation degree and a theoretical 20% methacrylation (O1M20A) degree were synthesized according to a similar method as described in the literature.^S1–2^ Briefly, 10 g of sodium alginate was dissolved in 900 mL of deionized water (diH_2_O) overnight, and 108 mg of sodium periodate (NaIO_4_) in 100 mL of diH_2_O was rapidly added to the alginate solution under stirring in the dark at room temperature (RT). After reaction for 24 h, 19.52 g of 2-ethanesulfonic acid (MES) and 17.53 g of sodium chloride (NaCl) were added, and the pH was adjusted to 6.5 with 5 N sodium hydroxide (NaOH). Then 1.18 g of *N*-hydroxysuccinimide (NHS) and 3.89 g of 1-ethyl-3-(3-dimethylaminopropyl)carbodiimide hydrochloride (EDC·HCl) were sequentially added to the mixture. After 10 min, 1.69 g of AEMA was added slowly. The solution was wrapped with aluminum foil to protect it from light and left to react for 24 h at RT. The mixture was then poured into 2 L of chilled acetone to precipitate out the crude OMA solid, which was further purified by dialysis against diH_2_O over 3 days (MWCO 3.5 kDa, Spectrum Laboratories Inc., Rancho Dominguez, CA). The dialyzed alginate solution was collected, treated with activated charcoal (0.5 mg/100 mL, 50-200 mesh, Fisher Scientific) for 30 min, filtered through a 0.22 μm filter and frozen at −80 °C overnight. The final O1M20A was obtained as white cotton-like solid through lyophilization for at least 10 days. The actual methacrylation of O1M20A was determined to be 6.1% from ^1^H NMR data according to the method described in the literature.^S3^ Note that the actual oxidation was not provided due to the overlap of the proton peak assigned to the CHO group (∼5.4 ppm) with the polymer proton peak (broad peak located at ∼5.1 ppm). The GelMA was the same material that was used in our previous work.^S4^ The ^1^H NMR spectra for newly synthesized OMA is shown in Figure S13.

#### 1.3 Cell expansion

Human mesenchymal stem cells (hMSC) were isolated according to the literature.^S5^ hMSC cells were expanded in GM supplemented with 10 ng/mL FGF-2 and NIH3T3 cells were expanded in GM. The culture was performed in a humidified incubator at 37 °C and 5% CO_2_ with medium changes every 2 or 3 days. The cells were harvested when they reached ∼80% confluence.

#### 1.4 OMA hydrogelation

Solution of OMA (3 w/v %) with or without GelMA (0.5 w/v %) in DMEM-LG containing photoinitiator (0.05 w/v %) was placed between two quartz plates with 1.0 mm spacers and UV crosslinked at ∼18 mW/cm^2^ for 45 s to form cell-free hydrogels, which were then ion printed using the ion-transfer printing (ITP) protocol described in Section 1.8 below and cut into specific geometries to culture in medium. For the mechanical property studies, dual-crosslinked OMA hydrogels were prepared by culturing the photocrosslinked hydrogels in a 0.5 M Ca^2+^ aqueous solution for 10 min. To encapsulate cells inside hydrogels, macromers in DMEM-LG containing cells were subjected to photocrosslinking ad described above, generating cell-laden hydrogels.

#### 1.5 Degradation and swelling tests

Photocrosslinked cell-free OMA hydrogels were fabricated as above (Section 1.4) and used for degradation and swelling tests. Circular hydrogel samples with a diameter of 8 mm (*d*_0_) were obtained via punching the bulk hydrogels using a biopsy punch. For the degradation test, the samples were frozen for 4 h at −80 °C and lyophilized for 2 days. The masses of the dried gels were measured as initial weights (*W*i). The dried hydrogels were rehydrated by culturing in 5 mL of GM under cell culture conditions (37 °C, 5% CO2), and the medium was changed every 3 days. At predetermined timepoints, the hydrogels were collected and dried by lyophilization to obtain dried mass (*Wd*). Degradation expressed as the mass loss was quantified as (*Wi*-*Wd*)/*Wi* × 100% (N = 3). For the swelling test, the samples were cultured for 4 h in 5 mL of H_2_O, PBS (pH 7.4), GM, or 0.5 M Ca^2+^ solution under RT. The diameters of the swollen hydrogels (*d_S_*) were measured. The swelling was calculated with the following equation: *ds*/*d_0_* (N = 3).

#### 1.6 Rheology

Dynamic rheological examination of the as-prepared and swollen/shrunk photocrosslinked cell-free OMA hydrogels was performed to measure the hydrogel storage moduli (G’) with a Kinexus ultra+ Highest specification rheometer (Malvern Panalytical, Malvern, United Kingdom). In oscillatory mode, a parallel plate geometry (8 mm diameter) measuring system was employed, and the gap was set to 1 mm. After each hydrogel was placed between the plates, all the tests were carried out at RT. Oscillatory frequency sweep (0.1∼100 Hz at 1 % strain) tests were performed to measure G’.

#### 1.7 Young’s modulus measurement

The elastic moduli of the as-prepared and swollen/shrunk photocrosslinked cell-free OMA hydrogels were determined by performing uniaxial, unconfined constant strain rate compression testing at RT using a constant crosshead speed of 0.8%/sec on a mechanical testing machine (225lbs Actuator, TestResources, MN, USA) equipped with a 5 N load cell. The Young’s modulus of each sample was determined using the first non-zero slope of the linear region of the stress-strain curve within 0 ∼ 10% strain (N = 3).

#### 1.8 ITP

ITP was used to create a crosslinking density gradient across the thickness of photocrosslinked OMA hydrogels. In a typical protocol, the filter paper (FisherBrand, qualitative P5, medium porosity, Fisher Scientific) was soaked in a Ca^2+^ bath at a concentration of 0.1 − 0.5 M (specified in experiments) for 30 min. The soaked filter paper was taken out of the ion bath and placed onto the surface of the photocrosslinked OMA hydrogel for 30 − 120 s (specified in experiments). The hydrogel with a Ca^2+^-crosslinking gradient was tailored into specific shapes (e.g., square, circle, and bars) and then cultured in medium for 10 min at RT. Hydrogel images were then taken for analyzing the shape changes. The bending angles were quantified according to a method previously described in the literature.^S6^

#### 1.9 Reprogramming and reversibility study

Reprogramming was realized through EDTA treatment and a subsequent re-ITPing. Briefly, ITPed hydrogels after deformation were transferred from the culture medium to 10 mL of EDTA solution (5 mM) in H_2_O (for cell-free hydrogels) or DMEM-LG without adding NaHCO_3_ (for cell-laden hydrogels), which were further cultured at RT (no shaking) or 37 °C under shaking (2.37 rev/s) (Bellco Glass 7744-01010 orbital shaker, Bellco Biotechnology, NJ, USA) for over 1.0 h to recover the shapes. Subsequently, the EDTA-treated samples were transferred into H_2_O (for cell-free hydrogels) or DMEM-LG (for cell-laden hydrogels) for 5 min to rinse away the residual EDTA within hydrogels. Then, the recovered hydrogels were reprogrammed via another ITP process on either the same side or the other side as described in Section 1.8, and this was followed by culture in medium to obtain another deformed shape. Reversibility was assessed by repeating the above process for five cycles.

### 2.0 Live/dead staining

The viability of cells was assessed using live/dead staining comprised of FDA and EB. The staining solution was freshly prepared by mixing 1 mL of FDA solution (1.5 mg/mL in DMSO) and 0.5 mL of EB solution (1 mg/mL in PBS) with 0.3 mL PBS (pH 8). 20 μL of staining solution per 1 mL of culture medium was added to sample medium and incubated for 5 min at RT. Fluorescence images of the samples were taken using a Nikon Eclipse TE300 fluorescence microscope (Nikon, Tokyo, Japan) equipped with a 14MP Aptina Color CMOS digital camera (AmScope, Irvine, CA).

#### 2.1 In vitro cartilage-like tissue engineering, biochemical quantification, and histological staining

The hMSCs at passage 5 (P5) were used for the chondrogenesis study. The cells were harvested for cell-only bioprinting when they reached ∼80% confluence. Square cell-laden hydrogels with a length of 18 mm were fabricated as above (Section 1.4). For cartilage-like tissue formation, the hMSC-laden hydrogels were cultured in 8 mL of chondrogenic medium in a humidified incubator at 37 °C with 5 % CO_2_ over a course of 21 days, and 4 mL of medium was changed every other day. The cartilage-like tissues were obtained at day 21 and used for biochemical analysis, histological staining, and ITP-induced shape programming and reprogramming. For controls, cell-laden hydrogels of identical dimensions, prepared under the same conditions, were cultured in cell growth medium and collected on day 21 for further investigation.

For biochemical analysis, the engineered cartilage tissues were first homogenized in 0.5 mL of papain buffer (Sigma) at 0 °C for 1 min and then digested in a total of 1.0 mL of papain solution (Sigma) at 65 °C for 24 hours and centrifuged for 10 min at 15,000 rpm, and then the supernatants were collected for DNA and glycosaminoglycan (GAG) quantifications (N = 3).

Per the manufacturer’s instructions, a Picogreen assay kit (Invitrogen) was used to quantify the DNA content in the supernatant. Fluorescence intensity of the dye-conjugated DNA solution was measured using a microplate reader with an excitation of 480 nm and emission of 520 nm.

The GAG content was quantified using a DMMB (1,9-dimethylmethylene blue) assay.^S7^ 40 μL of supernatant from the digested samples was transferred into 96-well plate, to which 125 μL of DMMB solution was then added. Absorbance at 595 nm was recorded on a microplate reader. GAG content was normalized to DNA content.

The cartilage-like tissues were fixed in 10% neutral buffered formalin (NBF) overnight at 4 °C, dehydrated, and embedded in paraffin. Briefly, tissue samples were cut into 5 μm thick sections using a Leica RM2255 rotary microtome (Leica Microsystem Ltd., Milton Keynes, UK). Slide sections were then deparaffinized and stained with Hematoxylin and Eosin (H&E) to observe gross cell and tissue morphology,^S8^ and Safranin O (SafO) with a Fast Green counterstain^S9^ and Toluidine Blue O (TBO) for glycosaminoglycan (GAG) indication.^S10^ Stained samples were imaged using a fluorescence microscope under bright field.

---

**Scheme S1.**
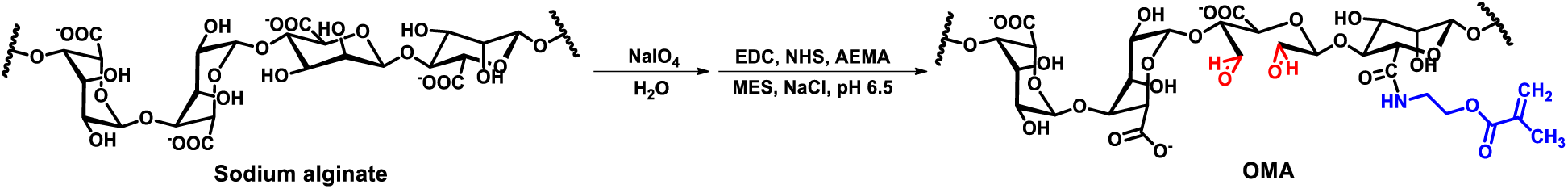
Synthesis of OMA.

**Figure S1.**
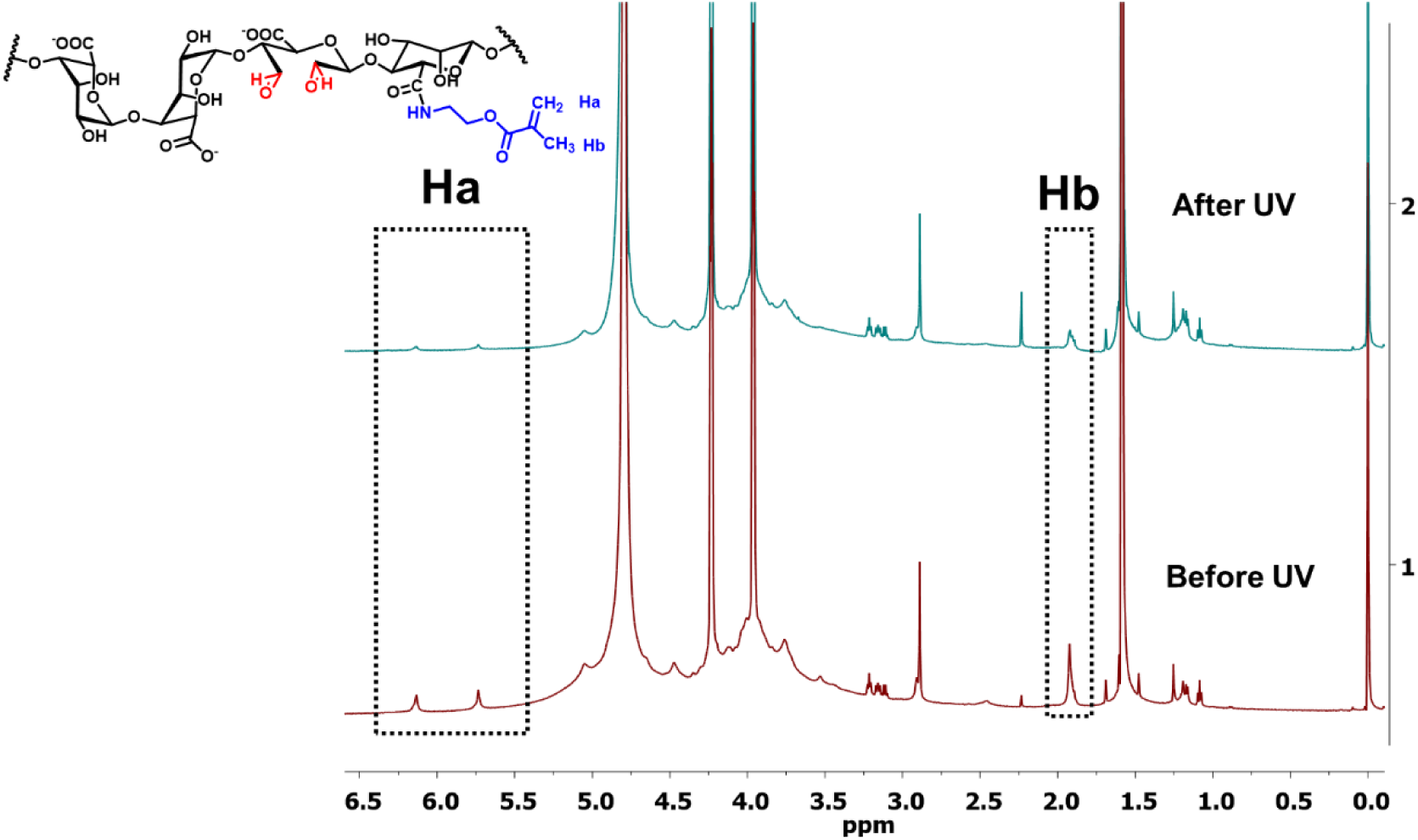
The weakening of the methacrylate protons signal on OMA macromers (3%, w/v) after 45 s UV irradiation in D_2_O with 0.05% (w/v) PI.

**Figure S2.**
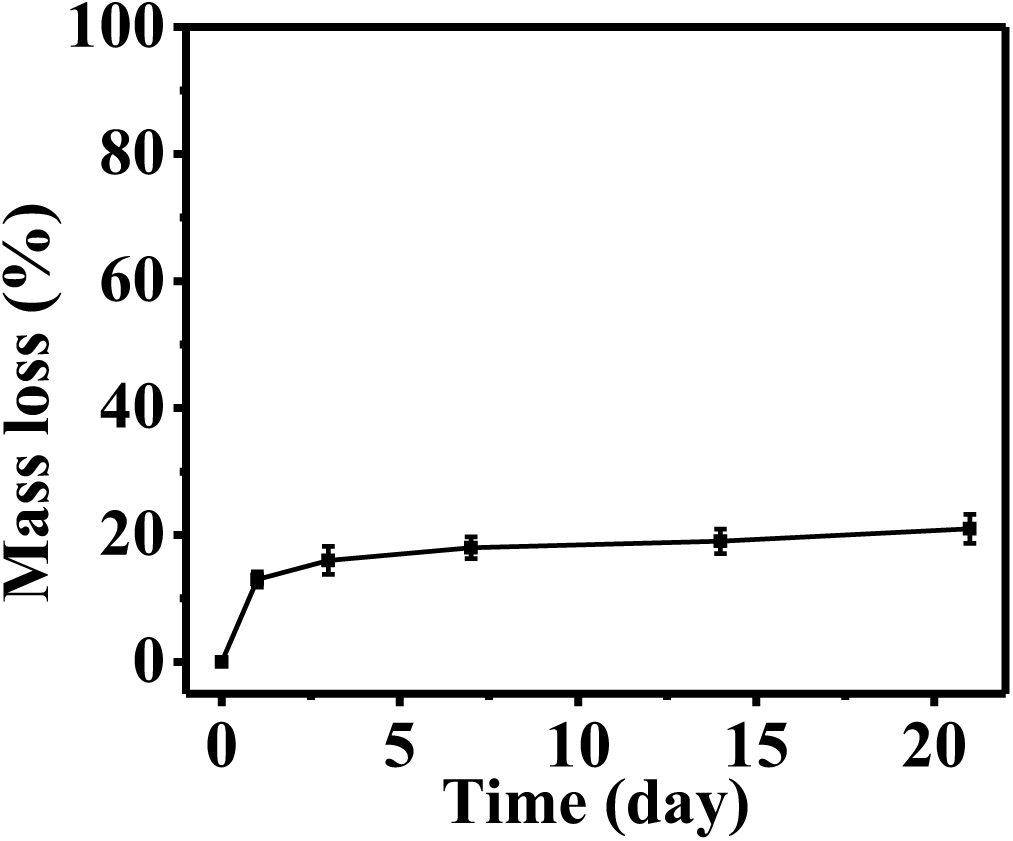
Degradation profile of photocrosslinked OMA hydrogel during culture in cell GM at 37 °C. *N* = 3, data are presented as mean ± SD.

**Figure S3.**
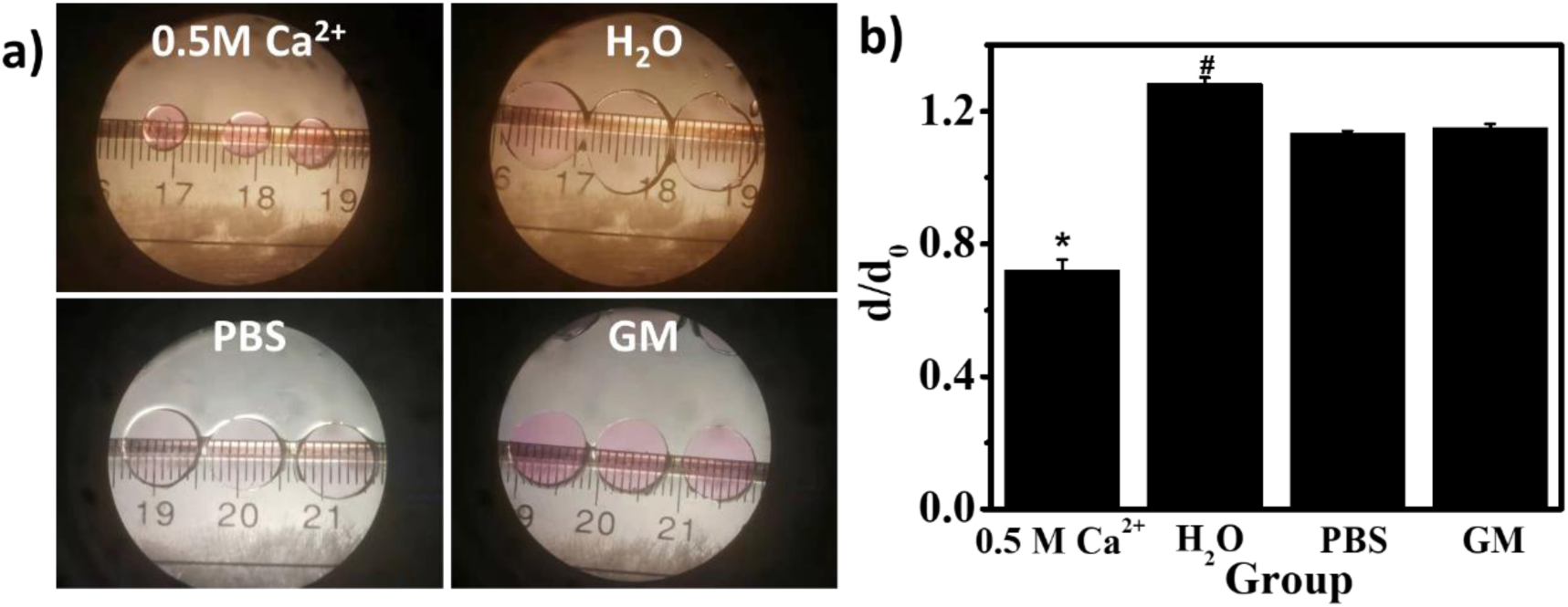
(a) Photomicrographs of three photocrosslinked OMA hydrogel discs after culture in different media for 30 min at room temperature. (b) Swelling/deswelling of photocrosslinked OMA hydrogel after culture in different media, where *d* is the diameter of hydrogel discs after culture and *d_0_* is the original hydrogel disc diameter (8 mm). *^,#^*p* < 0.05 compared to other groups. *N* = 3, data are presented as mean ± SD.

**Figure S4.**
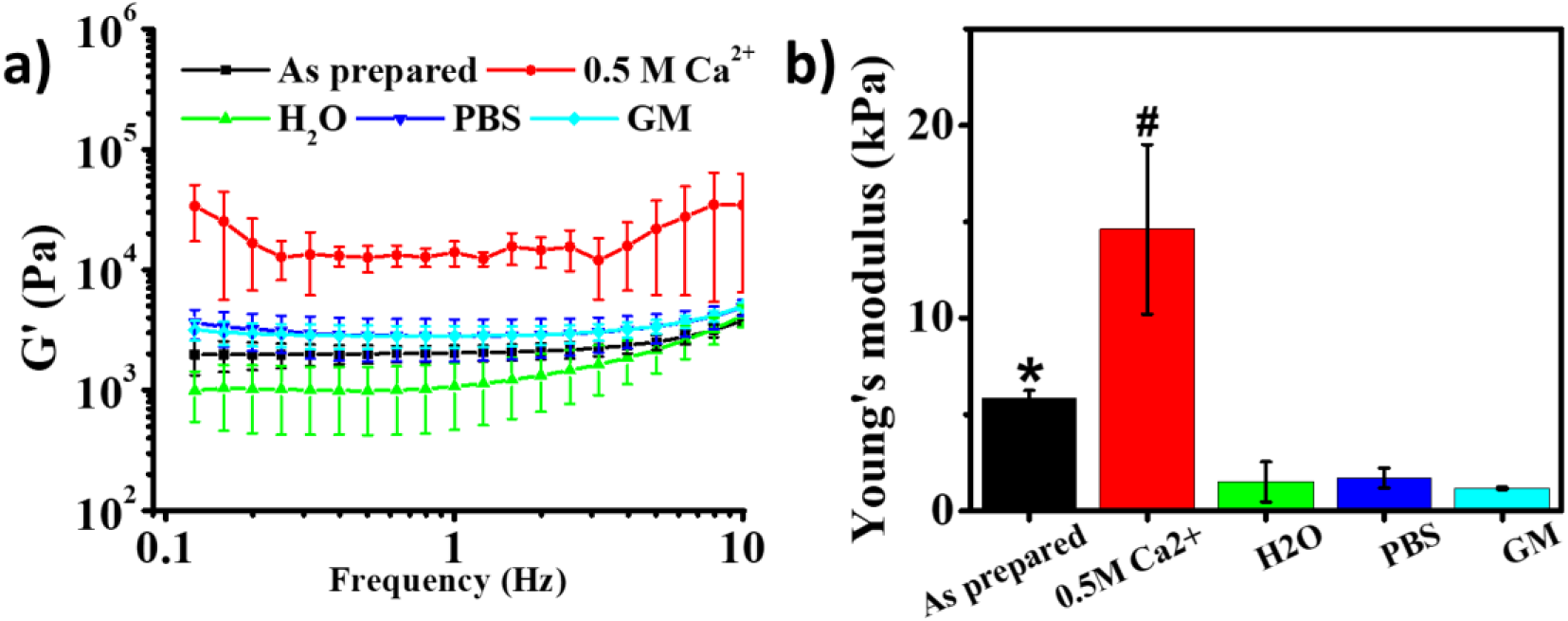
(a) Storage and (b) Young’s moduli of photocrosslinked OMA hydrogels after culture in different media for 30 min at room temperature. *^,#^*p* < 0.05 compared to other groups. *N* = 3, data are presented as mean ±SD.

**Scheme S2.**
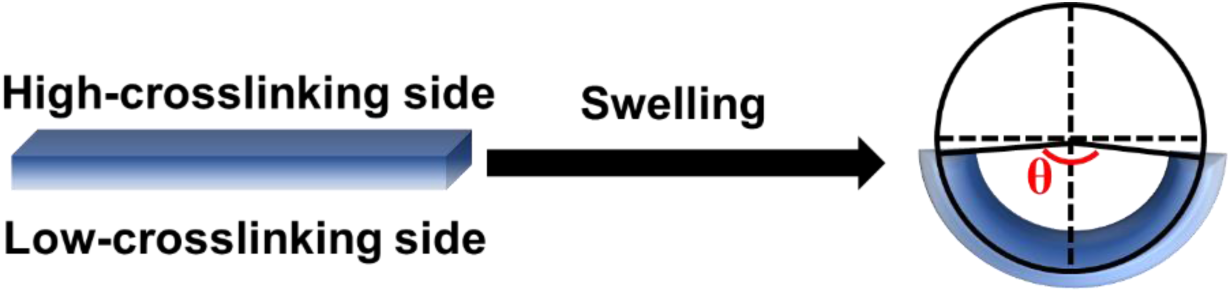
The bending of gradient OMA hydrogel slabs with high-crosslinking side facing inward and the definition of bending angle θ.

**Figure S5.**
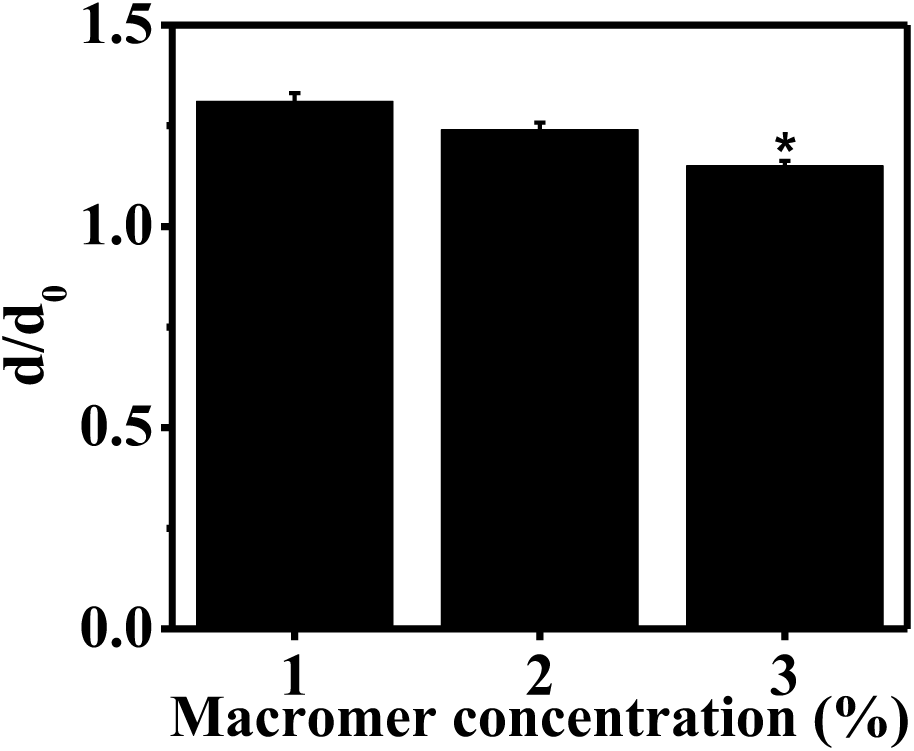
Swelling of OMA hydrogels versus macromer concentration in GM. \**p* < 0.05 compared to other groups. *N* = 3, data are presented as mean ±SD.

**Figure S6.**
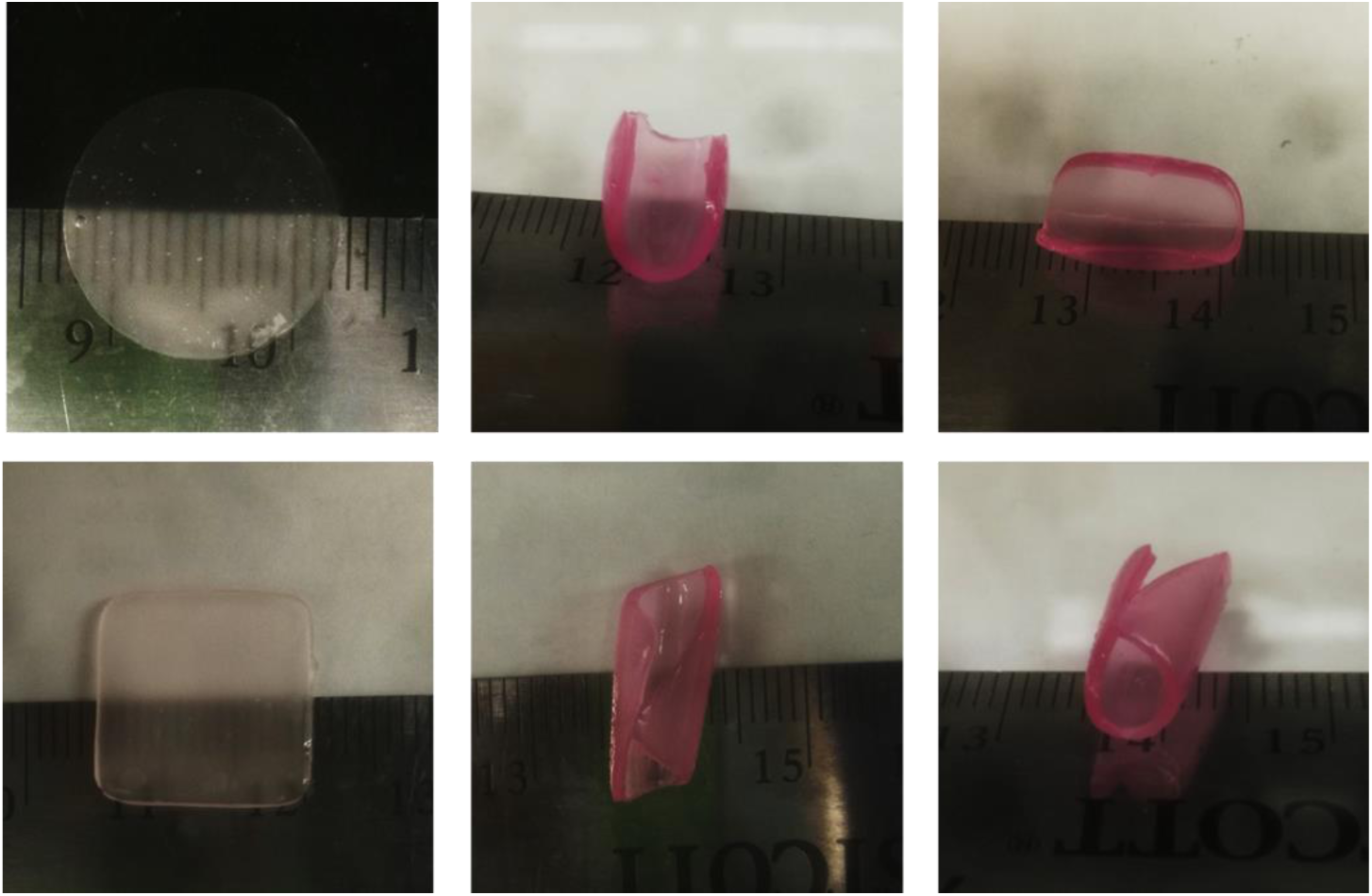
ITPed large cell-laden hydrogel disc and square hydrogel slab and their deformed shapes. Hydrogel disc: diameter 10 mm, square hydrogel slab: length 10 mm. ITP time: 5 min, culture in GM for 5 min.

**Figure S7.**
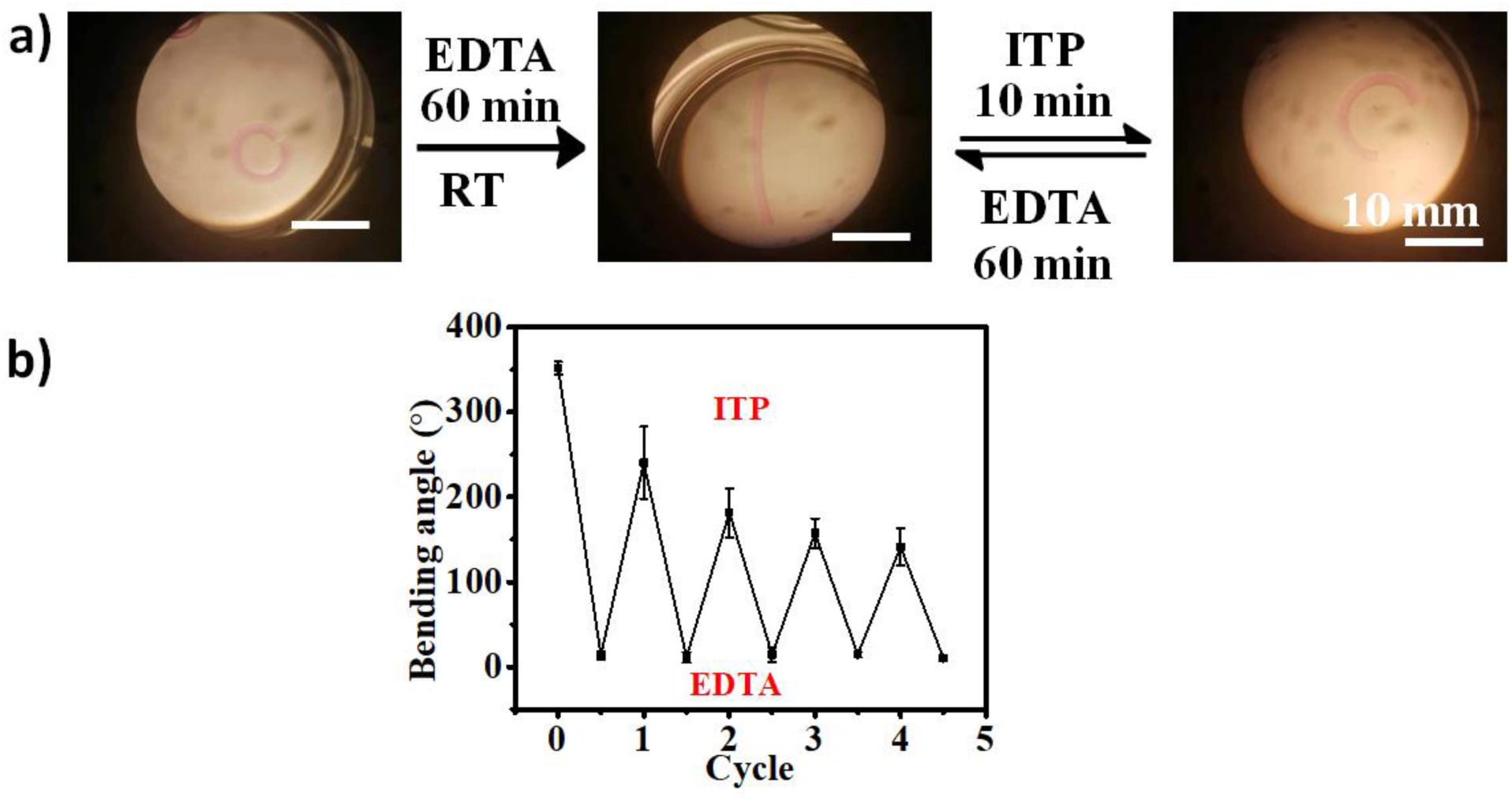
Demonstration of the reprogramming of cell-free hydrogel bars at RT. (a) Switch of a representative hydrogel bar between stretched and bent modes. (b) Quantitative bending angles of hydrogel bars at the two modes. H_2_O was used as medium for this study. *N* = 3, data are presented as mean ±SD.

**Figure S8.**
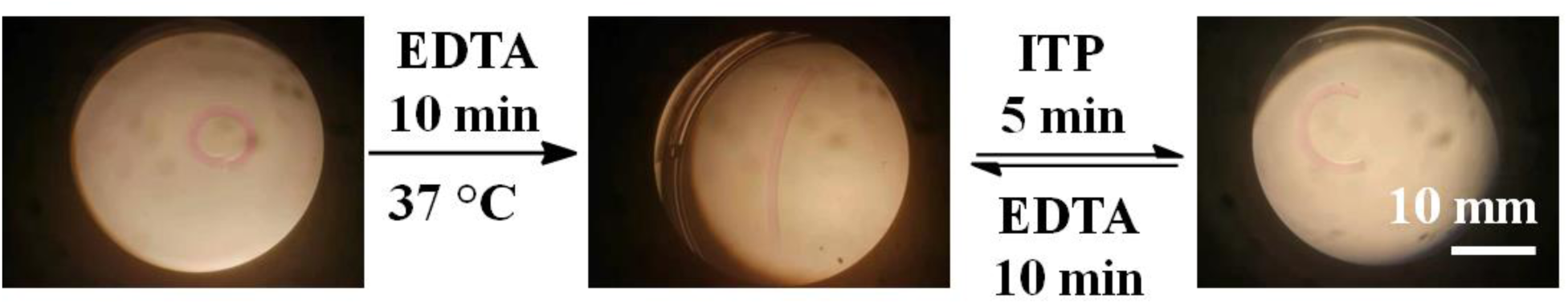
Switch of a representative hydrogel bar between stretched and bent modes at 37 °C. H_2_O was used as the solution for this study.

**Figure S10.**
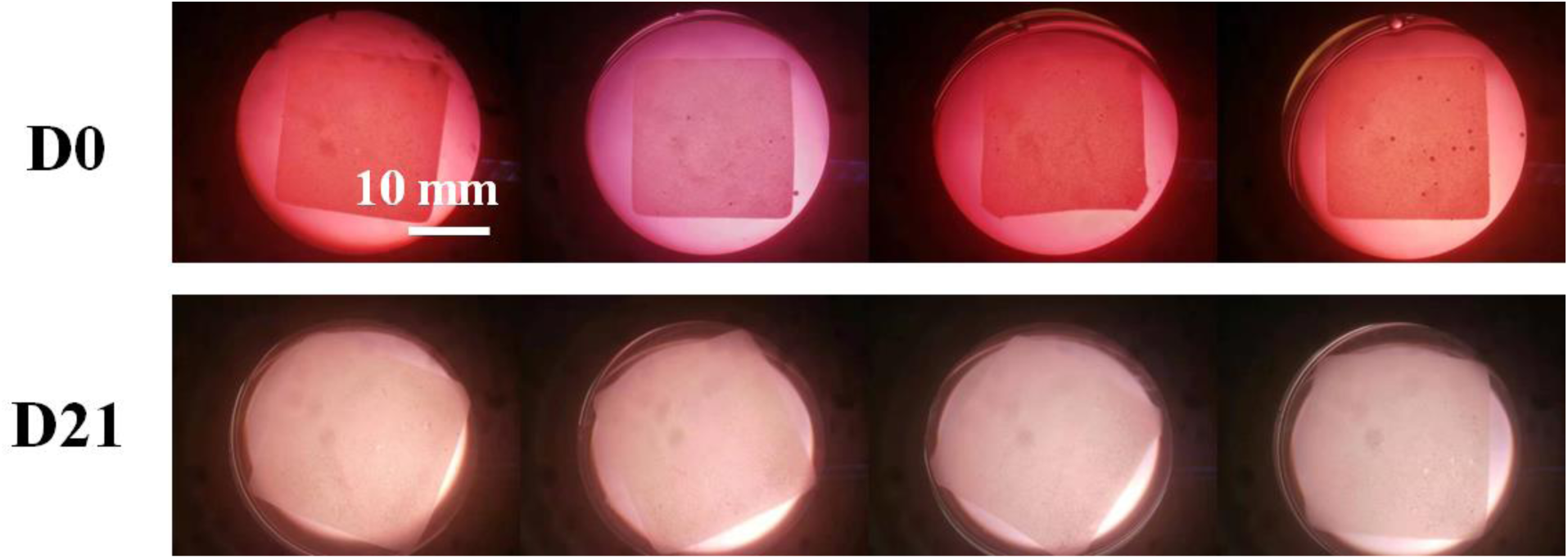
hMSC-laden bioconstructs (a) before and (b) after culture in chondrogenic medium for 21 days.

**Figure S11.**
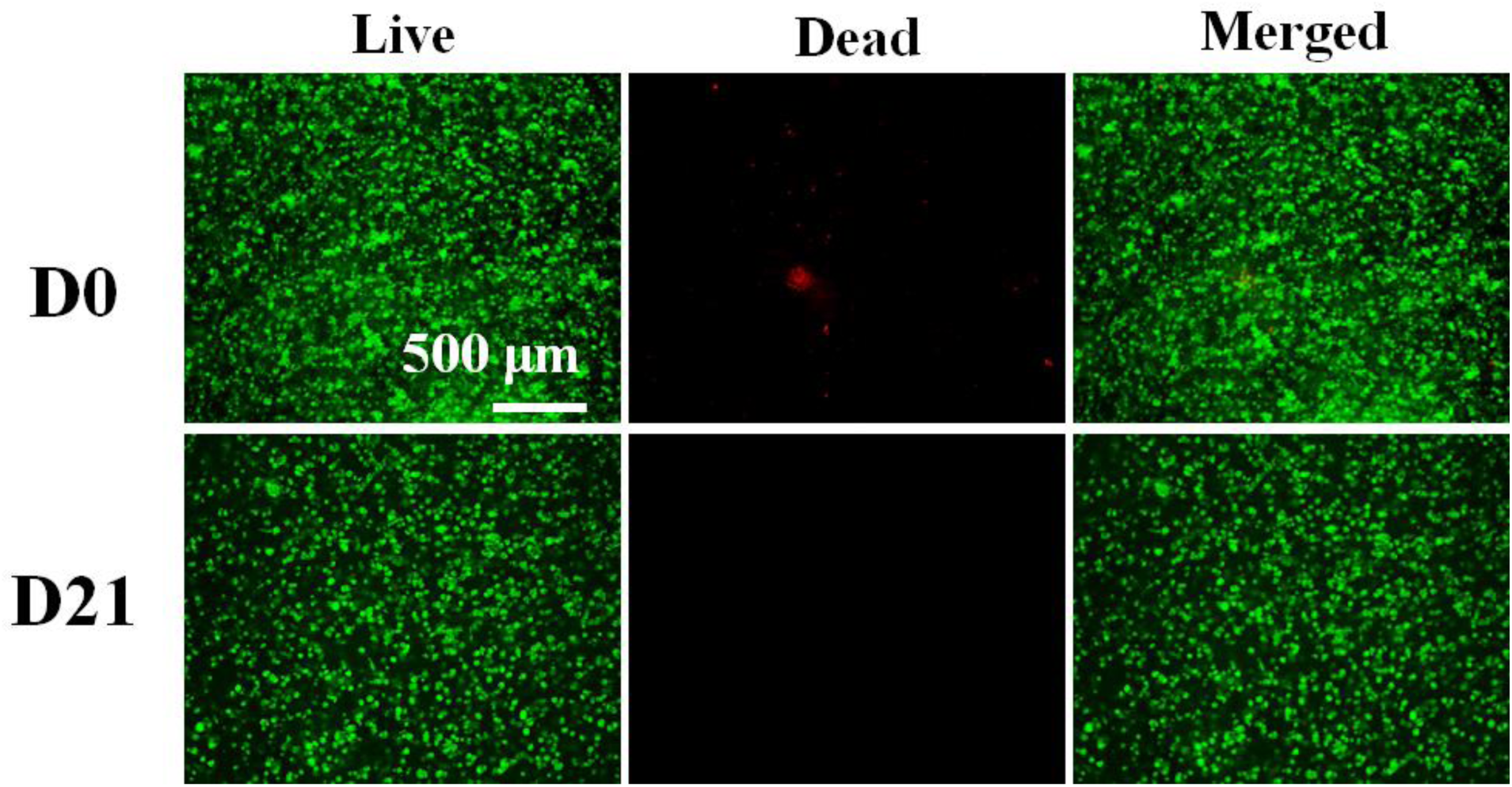
Live/dead staining of hMSC-laden hydrogels at D0 and D21.

**Figure S12.**
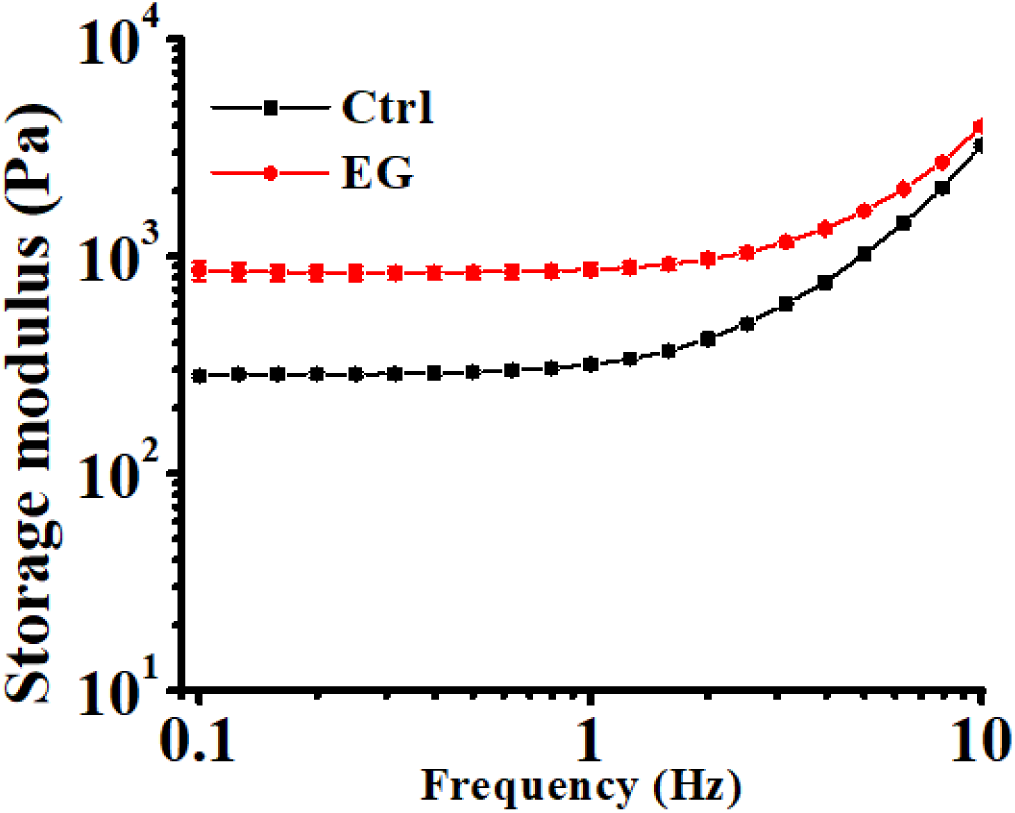
Storage modulus (G’) of bioconstructs from Exp and Ctrl groups. *N* = 3, data are presented as mean ±SD.

**Figure S13.**
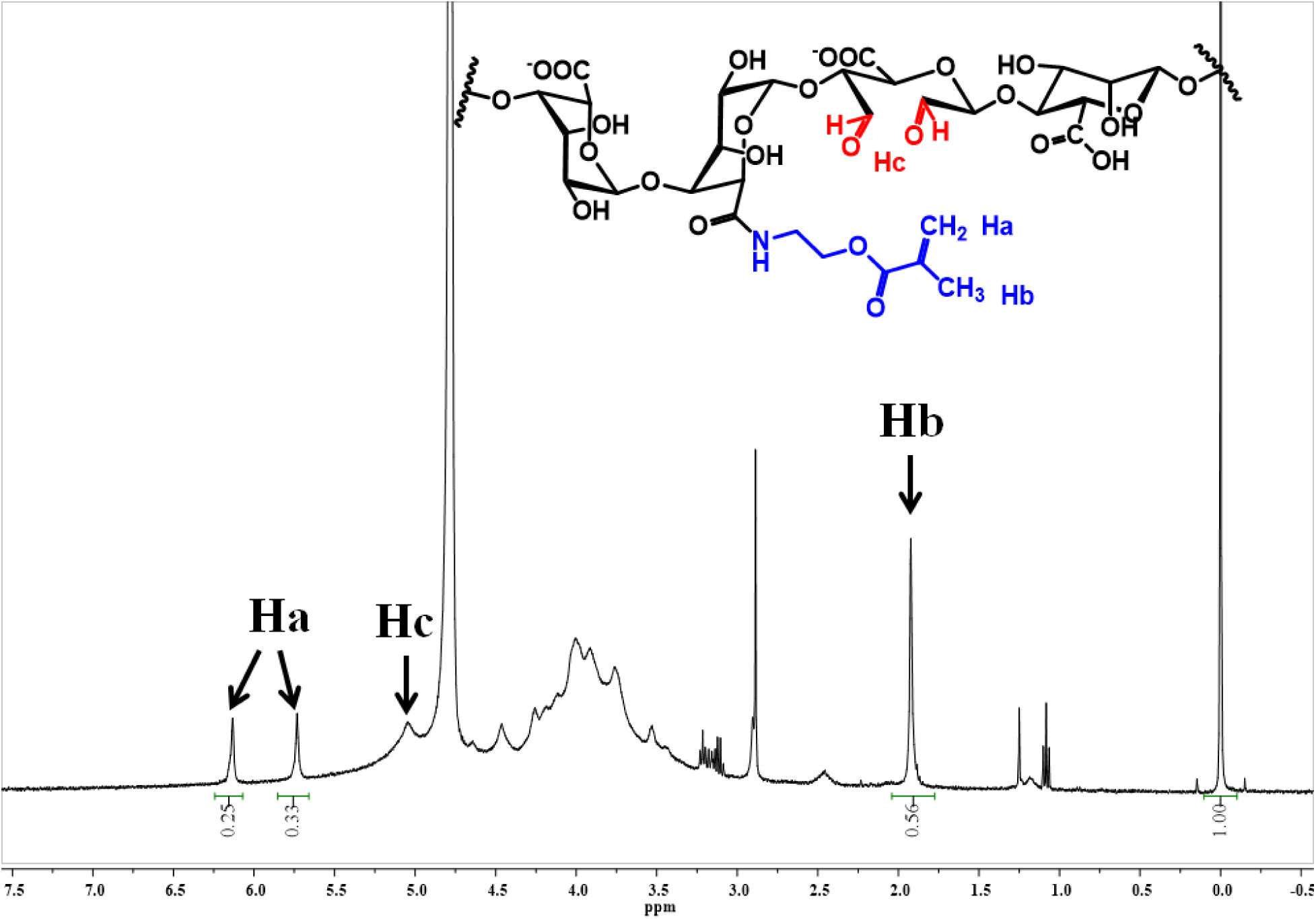
^1^H NMR spectrum of OMA (D_2_O, 2 w/v %).

